# TNF⍺-driven Aβ aggregation, synaptic dysfunction and hypermetabolism in human iPSC-derived cortical neurons

**DOI:** 10.1101/2024.08.30.610594

**Authors:** Alicia González Díaz, Elisa Belli, Benedetta Mannini, Gustavo Antonio Urrutia, Michele Vendruscolo

**Affiliations:** Centre for Misfolding Diseases, Yusuf Hamied Department of Chemistry, University of Cambridge, UK; WaveBreak Therapeutics, Cambridge, UK; Department of Experimental and Clinical Biomedical Sciences Mario Serio, University of Florence, Italy

## Abstract

Alzheimer’s disease (AD) patients exhibit an increased load of Aβ aggregates in the brain parenchyma. The neurotoxic nature of these aggregates has been underscored by recent advances in therapies aimed at reducing their load. To make further progress towards the development of increasingly effective treatments, there is a still largely unmet need for reliable cell models that comprehensively recapitulate aggregate-driven AD pathology. Here, we report a robust and scalable pipeline for generating human iPSC-derived cortical neurons that display Aβ aggregates in their axonal projections. This phenotype is caused by a repeated dosage of tumour necrosis factor-alpha (TNFα) to simulate the chronic inflammatory environment characteristic of AD and enhanced in neurons carrying the Swedish mutation. In association with the increased Aβ deposits in the cell bodies, this cell model exhibits other key hallmarks of AD, including structural alterations of synapses, electrophysiological asynchronous hyperactivity, and hypermetabolism. Overall, these results illustrate how repeated TNFα treatment models central aspects of AD pathology, and provides a platform that could be used for facilitating the translation of potential drugs to clinical applications.

## Introduction

Alzheimer’s disease (AD) represents a formidable challenge for our ageing populations (Nichols et al., 2022; Scheltens et al., 2021), even as disease-modifying therapies are becoming available (Cummings et al., 2024a; Cummings et al., 2024b; Sims et al., 2023; Van Dyck et al., 2023). At the histopathological level, AD is characterized by the progressive accumulation of Aβ into amyloid plaques (Hampel et al., 2021; Knopman et al., 2021; Selkoe and Hardy, 2016). This phenomenon may begin at early stages of the disease, triggering a cascade of pathological events, including synaptic dysfunction, mitochondrial impairment, inflammation, and oxidative stress, ultimately leading to the progressive loss of neurons. (Hampel *et al*., 2021; Knopman *et al*., 2021; Selkoe and Hardy, 2016). The deposition of Aβ aggregates in the brain is also one of the main criteria for the early diagnosis of AD (Jack Jr et al., 2024; Jack Jr et al., 2018). The significance of Aβ aggregation is underscored by the fact that the only three disease-modifying therapies approved for AD patients, aducanumab (Sevigny et al., 2016), lecanemab (Van Dyck *et al*., 2023) and donanemab (Sims *et al*., 2023), specifically target this process.

Despite the relevance of Aβ aggregation, it is still challenging to establish cell models derived from human induced pluripotent stem cells (hiPSCs) to reproduce this phenomenon (Arber et al., 2017; Choi et al., 2014; Israel et al., 2012; Lin et al., 2018; Oksanen et al., 2017; Park et al., 2018; Penney et al., 2020; Raja et al., 2016; Shi et al., 2012a; Shi et al., 2012b; Wang et al., 2024). Several of these studies rely on complex three-dimensional cultures or artificial overexpression of several familial AD variants. Other strategies are based on the exogenous administration of high concentrations of Aβ monomers and pre-formed aggregates to cells cultured in scalable formats (Bassil et al., 2021; González Díaz et al., 2024). While the latter approach enables the characterization of the biological activity of certain aggregate populations, they do not readily capture the complexity of endogenous Aβ aggregation by not fully representing the diverse species found in a disease cell environment. This limitation underscores the need for developing robust models that: (i) accurately recapitulate endogenous Aβ aggregation within a cellular context, capturing the diversity of aggregate species present in AD and their biological activity responsible for the dysfunction phenotypes, and (ii) are scalable and reproducible for high-throughput applications aimed at targeting Aβ aggregation.

To address this need, in this work we report a cell model based on hiPSC-derived cortical neurons carrying the Swedish mutation (APP^Swe^), which are subjected to a chronic treatment with tumour necrosis factor α (TNFα), a cytokine that plays an important role in the brain immune system (Brenner et al., 2015). The Swedish mutation (KM670/671NL) in the APP gene results in altered β-secretase cleavage site, leading to enhanced production of Aβ_42_, the 42-residue aggregation-prone form of the peptide (Haass et al., 1995), which is the major component of the amyloid deposits in the AD brain (Hampel *et al*., 2021; Knopman *et al*., 2021; Selkoe and Hardy, 2016). However, human neuronal models with this mutation only recapitulate a modest elevated production of monomeric Aβ over time, with no Aβ aggregation and concomitant dysfunction present without any stressor (Zhou et al., 2022). TNFα has been previously reported to enhance formation of Aβ aggregates on the cell (Whiten et al., 2020). Upon release by activated microglia (Gao et al., 2023), it has been found elevated in the blood and brain of AD patients, as well as detected in amyloid deposits (Serafini et al., 2024).

By combining the Swedish mutation with the repeated dosage of TNFα, we established a hiPSC-derived neuronal model that recapitulates the formation of endogenous Aβ aggregates in the cell body, without the need for exogenous addition of pre-formed Aβ aggregates. Furthermore, we observed in the model additional key hallmarks of AD, including an increased asynchronous electrical activity and an elevated metabolic rate indicative of hypermetabolism. To develop and increase the robustness of this model, we applied strategies to minimise the variability intrinsic of stem cell systems. We optimised a differentiation protocol to convert hiPSCs into batches of millions of well-characterised neuronal progenitor cells (NPCs) and cortical glutamatergic neurons, assessing each population for the expression of a panel of phenotype-specific markers. We set thresholds for each marker expression level and ensure that only NPCs and neurons meeting these quality standards were used. Overall, this optimized and well-characterized hiPSC-derived neuronal model, exhibiting key AD hallmarks including Aβ aggregation, synaptic and metabolic dysfunction, provides a platform for studying AD pathology and screening potential therapeutics.

## Results

### Establishment of a robust and scalable differentiation protocol for hiPSC-derived cortical neuron

To establish a cell system that can be scalable and reliable, we performed a series of quality controls throughout the differentiation from the hiPSCs to mature cortical neurons. We cultured in parallel two hiPSC lines obtained from Bioneer, the Swedish mutant and the wild-type isogenic control (Frederiksen et al., 2019), and included two clones from each line. All the lines were banked, and quality controlled as follows. The hiPSCs were routinely examined in terms of their morphology and density on the plate. Only round and well-defined colonies were kept and any irregular colony, without bright edges or with clearly differentiated cells, was manually removed before passaging on Matrigel. The colony density was maintained at no more than 80% confluence of the plate. The resulting cultures were assessed for their expression levels of OCT4, SOX2 and SSEA4 (**Figure S1A-C**). OCT4 and SOX2 are transcription factors essential for self-renewal of embryonic or induced pluripotent stem cells (ESCs or hiPSCs) (Zhang and Cui, 2014), whereas the stage-specific embryonic antigen-4, SSEA4, is a cell surface glycosphingolipid also enriched in undifferentiated, pluripotent stem cells (Itokazu and Yu, 2015). The expression of these three markers was quantified using both immunofluorescence and flow cytometry, with successful cultures showing expression in >90% of cells for all three markers. To confirm hiPSCs pluripotency, we also performed a trilineage differentiation test on each clone. Notably, all clones exhibited a similar expression of markers of mesoderm (NCAM and Brachyury, **Figure S1D**), endoderm (CXCR4 and SOX17, **Figure S1E**) and ectoderm (NESTIN and PAX6, **Figure S1F**).

After establishing high-quality hiPSC banks, we focused on developing a reproducible differentiation protocol to convert hiPSCs into NPCs, and NPCs into neurons. To achieve this goal, we recognized the importance of controlling the nature, phenotype, and potency of the generated NPCs. NPC populations in the brain are heterogeneous and multipotent, with each type capable of producing a limited range of cell types (Strano et al., 2020). The molecular profile of NPCs, especially their transcription factor expression (Strano *et al*., 2020), determines their differentiation potential. Our goal was to generate cortical NPCs that could give rise to TBR1-positive cortical glutamatergic neurons, which are affected in early stages of AD. A recent comprehensive study reported that NPCs enriched in EMX1 and defective for NKX2.1, convert into such cortical excitatory neurons (Strano *et al*., 2020). Given this evidence, we tested, in parallel, three chemical differentiation protocols to identify the one that could generate EMX1-enriched NPC populations, and derived TBR1-positive neurons with higher reproducibility.

The first protocol (*Protocol 1*) (Shi *et al*., 2012a) employed a double SMAD inhibition strategy, involving the use of the small molecules SB431542 (to inhibit TGF-β signalling) and LDN193189 or Noggin to inhibit BMP signalling (Chambers et al., 2009). This step promotes neural induction by suppressing mesoderm and endoderm fates (Chambers *et al*., 2009). Subsequently, the protocol incorporated fibroblast growth factor 2 (FGF2) to induce the formation of neural rosettes. FGF2 is crucial in promoting the proliferation and survival of neural progenitor cells, as well as in patterning the developing neural tube (Zhang et al., 2001).

The second protocol (*Protocol 2*) (Shi *et al*., 2012a) was identical to *Protocol 1*, but used the WNT inhibitor CHIR99021 during the later stages of rosette development (days 12-17). WNT signaling plays a complex role in neurodevelopment, and its inhibition can promote the specification of certain neural subtypes, particularly those of the forebrain (Maroof et al., 2013). Lastly, the third protocol (*Protocol 3*) (Cao et al., 2017), implemented the use of cyclopamine, a Sonic Hedgehog (SHH) pathway inhibitor, from day 3 to day 9 of the differentiation process. SHH signaling is a key regulator of ventral-dorsal patterning (Ericson et al., 1995) in the developing neural tube (Gaspard et al., 2008). By inhibiting this pathway during the early stages of neural induction, this protocol aims to promote the generation of dorsal neural progenitors, that may give rise to cortical neurons (Cao *et al*., 2017). All these three protocols promote the stepwise differentiation of hiPSCs into neuroepithelia, primitive rosettes (structures that mimic the early neural tube), and mature rosettes (structures that more closely resemble the ventricular zone of the developing cortex), preceding the generation of NPCs (**Figure 1A**).

**Figure 1.**
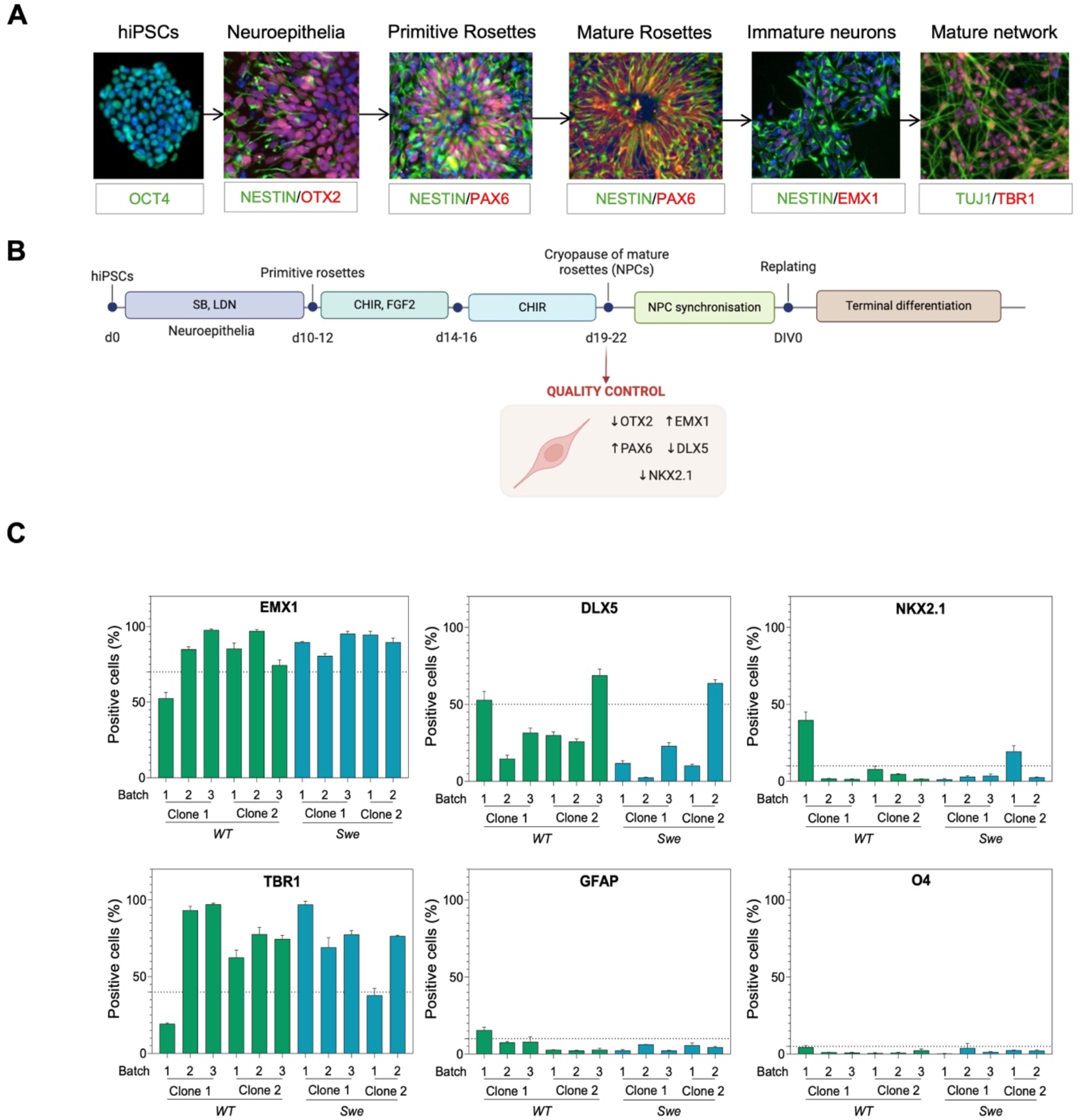
Illustration of the protocol to differentiate hiPSCs to a population of progenitors enriched in cortical markers. **(A)** Representative images of the key stages of differentiation from hiPSCs to mature cortical neurons and their respective ICC markers. **(B)** Schematic representation of the differentiation protocol. hiPSCs were induced to generate neuroepithelia by exposing them to 10 µM SB431542 (SB) and 100 nM LDN-193189 (LDN) for 10 to 12 days, until the appearance of primitive rosettes. At this point the media was supplemented instead with 1 µg/mL of CHIR-99021 (CHIR) and 20 ng/mL of FGF2 for 2 days to induce the expansion of NPCs. Subsequently, FGF2 was removed, and the rosettes were cultured to maturation until d19-d22, when the cultures were harvested for cryopreservation. Batches of NPCs were subjected to quality control (QC) by culturing them for 3-5 days for synchronization, and then replated for terminal differentiation (DIV0). **(C)** Markers of NPC and neuronal differentiation (TBR1, EMX1, DLX5) and culture purity (GFAP and O4). Quantification of immunofluorescence staining of neuronal cultures at DIV7 across several clones and NPC batches of wild-type and APP^Swe^ lines. In **C**, triplicate data are represented as mean ±SE.

To monitor the effectiveness of each protocol to convert hiPSCs to cortical NPCs, we followed the expression of the transcription factor OTX2, the proliferation marker Ki67 (Uxa et al., 2021), together with several dorsal (PAX6 (Elsen et al., 2018; Englund et al., 2005), EMX1 (Strano *et al*., 2020)) and ventral (DLX5 and NKX2.1 (Strano *et al*., 2020)) progenitor markers, at three different time points: d14, d18 and d24 (**Figure S2A**). The protocols were originally tested in one clone per hiPSC line (wild-type and Swedish).

*Protocol 1* and *Protocol 2* progenitors exhibited the progression of OTX2 expression described during embryonic development, that is OTX2 levels increase during rosette formation (d14-d18) but are comparatively reduced at later stages (d24) (Beby and Lamonerie, 2013; Larsen et al., 2010; Zhang et al., 2018), when neuronal progenitor cells are mature, suggesting that such progenitors may not hold ventral features (**Figure S2A**). For both protocols, EMX1 and PAX6 (dorso-cortical progenitor markers) exhibited increased expression, contrarily to DLX5 and NXK2.1 (ventral progenitor markers), which decreased in expression over time (**Figure S2A**). A decrease in Ki67 marker indicated that an increasing proportion of cells were exiting the cell cycle and committing to a neuronal fate (**Figure S2A**). On the other hand, *Protocol 3* yielded the opposite results. Generated progenitors showed a lower early expression of OTX2, higher expression of DLX5 and NXK2.1 and a lower expression of EMX1 and PAX6. Therefore, this protocol was discarded since it was not generating progenitors enriched in dorso-cortical markers.

In parallel to protocol exploration, to optimize the working capacity and ensure consistency across experiments, we scaled up the differentiation formats from multi-well plates (such as 6-well plates) to larger T75 and T175 flasks. This expansion significantly increased the yield of neural progenitor cells (NPCs), enabling us to conduct a broader range of experiments with well-characterized, high-quality, NPC cultures. In addition, we introduced a cryopause step into our workflow, allowing us to freeze large batches of NPCs after 19 to 22 days of differentiation (see methods). A freezing step makes it possible to pause the differentiation and to maintain large batches of cells at the stage of neuron progenitor cells (NPCs), aiding the consistency and throughput required for drug testing.

Cryopreserved NPCs generated by *Protocols 1* and *2* were thawed and differentiated to neurons, following the protocols reported in the corresponding studies. Importantly, neurons were differentiated and cultured in microplates (either 96 or 384), to allow for further high-throughput screening applications. At 10 days *in vitro* (DIV10), the levels of expression of the cortical marker TBR1 were examined by immunocytochemistry. TBR1 is a transcription factor crucial for the development of the cerebral cortex, particularly in the specification of early-born neurons excitatory glutamatergic neurons of the layer VI and subplate (Bedogni et al., 2010; Han et al., 2011; Hevner et al., 2001). *Protocol 2* yielded a higher percentage of TBR1^+^ neurons both for the wild-type and Swedish clones (**Figure S2B**,**C**) as compared to *Protocol 1*.

Therefore, we finally selected *Protocol 2* (depicted in **Figure 1B**) and tested how reproducible it was to generate cortical NPCs and derived TBR1^+^, Ki67^-^ post-mitotic cortical neurons (**Figure S2D**). For this purpose, we differentiated the wild-type and Swedish clones, three times each, from hiPSCs to NPCs, which were cryopreserved. NPCs were then thawed and immunoassayed for EMX1, PAX6, DLX5, NKX2.1. Cells were differentiated to neurons and stained for the astrocytic marker GFAP and the oligodendrocyte specific marker O4, to ensure that we were generating cultures without contaminating, non-desired phenotypes (**Figure 1C**).

Based on previous findings, we established stringent quality control criteria for our neural progenitor cell (NPC) batches (Strano *et al*., 2020). These criteria included expression thresholds of >70% for EMX1, < 50% for DLX5, and < 20% for NKX2.1, analysed as the percentage of positive nuclei over the total nuclei population.

Most NPC batches derived from various clones and lines met these criteria, showing appropriate enrichment of EMX1 and low expression of DLX5 and NKX2.1. However, we identified certain batches that failed to meet our quality standards (**Figure 1C**). For instance, Batch 1 from Clone 1 of wild-type hiPSCs showed only 50% EMX1 expression, with the remaining cells expressing NKX2.1 (**Figure 1C**). Similarly, Batch 1 of Clone 2 from Swedish hiPSCs exhibited NKX2.1 expression exceeding our 20% threshold (**Figure 1C**). Importantly, when these suboptimal NPC batches were differentiated into neurons, they demonstrated lower expression of TBR1 at DIV7 (**Figure 1C**), falling below our established threshold of 50%. The batch derived from wild-type Clone 1 also showed a slightly higher presence of GFAP-positive cells, indicating reduced purity of the neuronal culture.

The same NPCs were differentiated to cortical neurons, and the levels of TBR1 expression were tracked to ensure consistent expression patterns throughout neuronal maturation (DIV10, DIV17, DIV22) (**Figure S2D**). This longitudinal analysis confirmed that NPC batches failing our initial quality control criteria consistently produced neurons with lower TBR1 expression (**Figure 1C** and **S2D**).

In summary, our optimized differentiation protocol and rigorous quality control measures ensure the generation of well-characterized cortical neuron populations. This robust and scalable system provides a solid foundation for studying AD pathology, aiming to offer enhanced reliability and reproducibility.

### Chronic treatment with TNFα promotes the formation of Aβ aggregates in the axonal projections of APP^Swe^ cortical neurons

To develop a model of AD, first we aimed to recapitulate Aβ aggregation in the neurons. A previous report showed that a human neuronal model carrying the Swedish mutation exhibits a modest elevation of monomeric Aβ over time, with no Aβ aggregation and concomitant dysfunction (Zhou *et al*., 2022), suggesting that the presence of this mutation does not readily trigger an Aβ aggregation phenotype. We then adopted the strategy of further stressing the system by applying a chronic treatment with 20 ng/mL of TNFα from DIV14 to DIV41. As a readout of Aβ aggregation, we quantified the levels of WO2-positive puncta in the neurons over time. Notably, we observed that the puncta were localised in the axons and that after DIV29 only neurons treated with TNFα demonstrated a significant increase in the puncta levels normalised by neuronal area (**Figure 2A**,**B**). Given that WO2 binds to both APP and Aβ, we explored a panel of antibodies reported to detect only APP (LN27), both APP and Aβ (6E10), both Aβ_40_ and Aβ_42_ isoforms (3A1) and only Aβ_42_ (12F4) (**Figure 2C,D**). APP or Aβ-specific antibodies showed different staining patterns on the cultured neurons (**Figure 2C**). While most of the LN27 signal was spread across the cell body and perinuclear areas, the 3A1 and 12F4 signals were spotty and mostly localised in the axonal projections. The APP and Aβ antibody 6E10 showed a mixed staining pattern, with both spread and clustered signal in the neuronal body and axons, respectively. We then quantified the levels of spots that stained positive for all tested antibodies in the neuronal projections (**Figure 2D**). For this quantification, we excluded spots that co-stained with DAPI, which usually derived from cellular debris. Independently of the epitope-specificity of the antibody used in the testing, the levels of puncta demonstrated a significant increase after at least two weeks of chronic treatment with TNFα.

**Figure 2.**
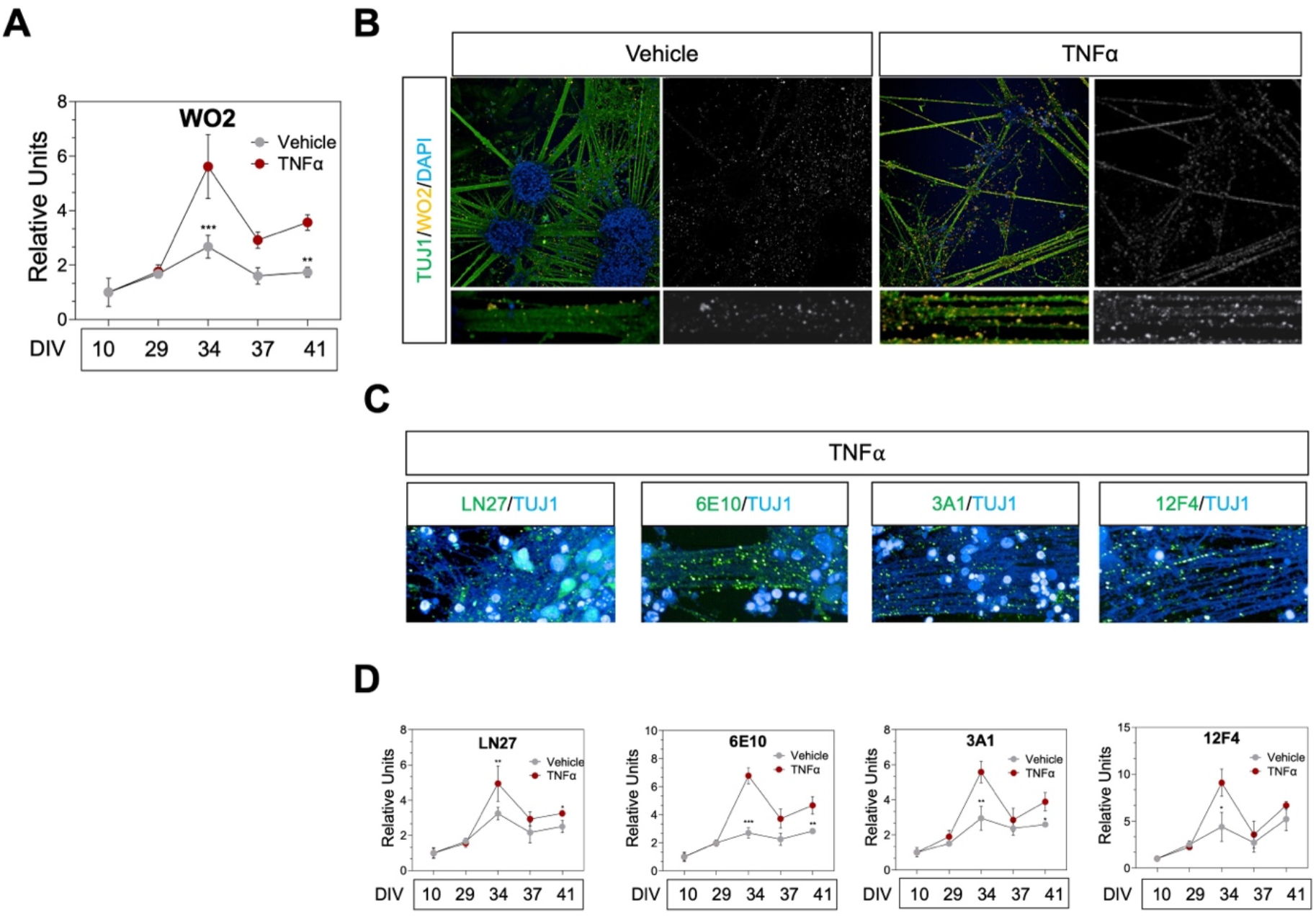
Chronic treatment with TNFα increases Aβ puncta levels in APP^Swe^ cortical neurons. **(A)** Longitudinal quantification of Aβ puncta in neuronal projections with the anti-APP/Aβ antibody WO2. Differentiated APP^Swe^ neurons were cultured in the presence of 20 ng/mL of TNFα or vehicle (D-PBS). WO2-positive puncta were quantified within TUJ1-positive projections. **(B)** Representative images at DIV34 of APP^Swe^ neurons treated with vehicle or TNFα. Coloured panels show the staining of nuclei with DAPI (blue), projections with TUJ1 (green) and Aβ inclusions with WO2 (orange). Black and white panels depict the WO2 channel. Zoomed-in sections of the neuronal projections are shown in the bottom panels. **(C)** Representative images of APP^Swe^ neurons stained with a panel of antibodies that recognize APP (LN27), both APP and Aβ (6E10), both Aβ_40_ and Aβ_42_ isoforms (3A1) and only Aβ_42_ (12F4) (**D)** Quantification of the APP/Aβ puncta detected with a panel of anti-APP/Aβ antibodies. Puncta were quantified within TUJ1-positive axonal projections of APP^Swe^ neurons in the presence of TNFα compared to vehicle controls. In **A, C** and **D**, triplicate data were normalized to DIV10 and are presented as mean ± SE. Two-way ANOVA was carried out to identify differences between the TNFα and vehicle treated samples. Significance levels are as follows: (^*^) *P* < 0.05, (^**^) *P* < 0.01, (^***^) *P* < 0.001.

### TNFα-driven accumulation of Aβ correlates with synaptic and metabolic dysfunction in APP^Swe^ cortical neurons

We then assessed whether the increase in the levels of Aβ puncta in TNFα-treated neurons correlated with alterations in synapses, network activity and mitochondrial metabolism. We first measured the level of co-clustered SYN1 (a pre-synaptic marker) and PSD-95 (a post-synaptic marker) signal. Apposition of both markers is indicative of structural assembly of synaptic elements (**Figure 3A-C**). Chronic TNFα treatment decreased co-localisation of SYN1 and PSD-95 after DIV37 (**Figure 3C**). Similarly, lower levels of neurogranin and HOMER1, both post-synaptic markers which expression and distribution is altered in AD brains (Kvartsberg et al., 2015; Reddy et al., 2005; Urdánoz-Casado et al., 2021), were detected in TNFα treated neurons with more pronounced differences compared to the vehicle after DIV37 for both markers (**Figure 3C**). These alterations in the synaptic proteins seem to follow the appearance of the peak of Aβ deposition in axonal projections at DIV34.

**Figure 3.**
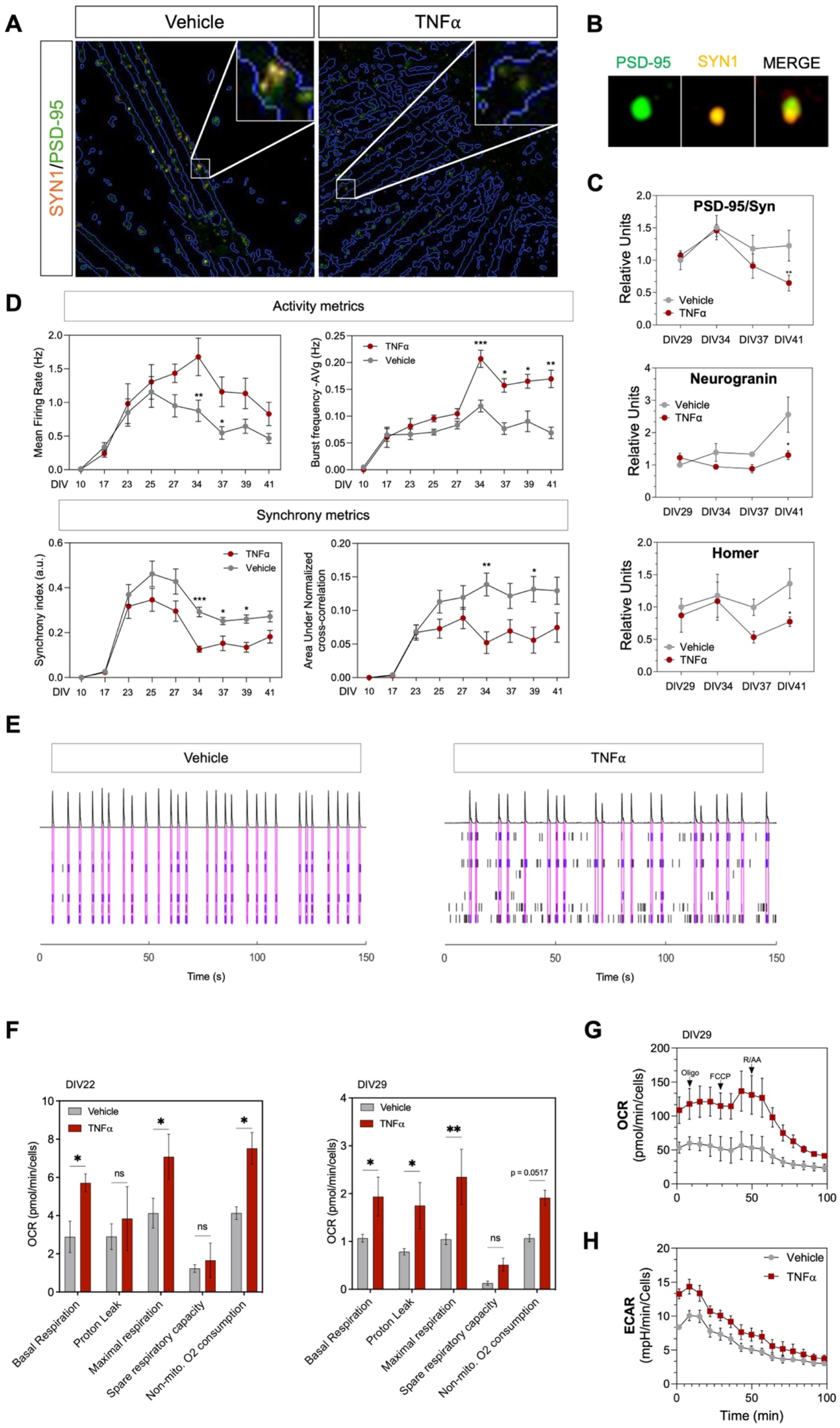
Chronic treatment with TNFα triggers synaptic dysfunction and hypermetabolism in APP^Swe^ cortical neurons. **(A)** Representative images of APP^Swe^ cortical neurons ± 20 ng/mL TNFα at DIV41, stained with anti-SYN1 (orange) and anti-PSD-95 (green) antibodies. Zoomed-in sections of axonal projections are shown on the top-right of each panel. **(B)** Representative pictures of the apposition (merge) of PSD-95 and SYN1, which is indicative of structural assembly of synaptic elements. **(C)** Longitudinal quantification of neurotransmitter ICC data from APP^Swe^ cortical neurons ±TNFα. Each marker was quantified only within TUJ1-positive projections. **(D)** Longitudinal quantification of metrics of activity (mean firing rate and Burst-Frequency average) and synchrony (synchrony index and area under normalized cross-correlation) and **(E)** raster plots derived from MEA wells containing APP^Swe^ cortical neurons ± 20 ng/mL TNFα. **(F)** Bar graphs showing the oxygen consumption rate (OCR) metrics at DIV22 and DIV29 under vehicle and TNFα treatments, including basal respiration, proton leak, maximal respiration, spare respiratory capacity, and non-mitochondrial oxygen consumption. **(G-H)** Line graphs depicting OCR and extracellular acidification rate (ECAR) over time for vehicle and TNFα-treated APP^Swe^ neurons. In **G** are indicated the time points of application of oligomycin (Oligo, ATP synthase inhibitor), FCCP (mitochondrial uncoupler), and rotenone/antimycin A (R/AA, inhibitors of the mitochondrial respiratory chain complex I and III, respectively). In **C, D, F, G** and **H**, triplicate data were normalized to the earliest DIV and are presented as mean ±SE. Two-way ANOVA was carried out to identify differences between the TNFα and vehicle treated samples within time points. Significance levels are as follows: (^*^) *P* < 0.05, (^**^) *P* < 0.01, (^***^) *P* < 0.001.

To correlate these observations with synaptic function, we measured the electrophysiological activity of neurons treated with vehicle or TNFα using microelectrode array (MEA) plates (**Figure 3D,E**). We observed opposite effects on neuronal activity and synchrony metrics. TNFα-treated neurons displayed a higher mean firing rate and more frequent bursts as compared to the vehicle (**Figure 3D**). By contrast, synchrony parameters such as the synchrony index or the area under normalised cross-correlation were significantly lower at late stages (DIV34 onwards) upon TNFα treatment (**Figure 3D**). More pronounced differences were again observed around DIV34 for all four metrics. These results suggest that APP^Swe^ neurons under TNFα stress show an association between asynchronous electrical activity and increased cellular Aβ cluster density.

Mitochondrial hypermetabolism has been reported to be an early event in the AD cascade, preceding synaptic dysfunction and cognitive decline in mice models and human patients (Bakhtiari et al., 2024; Naia et al., 2023). We thus set out to examine the effects of TNFα-driven dyshomeostasis on key parameters of mitochondrial respiration (**Figure 3F-H**). At DIV22 and DIV29, we measured the oxygen consumption rate (OCR) in basal conditions, and after treating the cells subsequently with oligomycin, carbonyl cyanide-4-(trifluoromethoxy)phenylhydrazone (FCCP) and rotenone/antimycin A. These compounds are used to inform on mitochondrial function.

For both DIV22 and DIV29, oxygen consumption of basal respiration, used to meet ATP synthesis, was significantly higher for TNFα-treated neurons, as shown by the higher levels of both OCR and extracellular acidification rate (ECAR) (**Figure 3G-H**). When measuring the basal respiration after inhibiting ATP synthetase with oligomycin, we observed higher levels of proton leak in neurons treated with TNFα, particularly at DIV29 (**Figure 3F**). Notably, proton leak is reported as a sign of mitochondrial damage (Gu et al., 2021). We then added FCCP to dissipate the proton gradient and measure the maximal respiration levels, observing that those were significantly higher for TNFα-treated neurons (**Figure 3F**). We also measured the spare respiration, defined as the difference between maximal and basal respiration (Gu *et al*., 2021). This parameter reports on the ability of the neurons to adapt to shifts in energy requirements. No significant differences were observed in the spare respiration between TNFα and vehicle-treated cells at both DIV22 and DIV29 (**Figure 3F**). By adding rotenone and antimycin A, we disrupted the electron transport chain. This procedure makes it possible to measure the non-mitochondrial respiration, referring to the levels of oxygen consumption from cellular sources other than mitochondria (Gu *et al*., 2021). We found that the metabolism of neurons treated with TNFα seems to depend less on mitochondrial oxygen production as compared to vehicle-treated cells (**Figure 3F**), suggesting that mitochondria may be impaired upon neuroinflammatory stress.

## Discussion

We reported a robust and scalable protocol to differentiate hiPSCs into cortical neurons to model key hallmarks of AD by using chemically defined factors. We used chemical differentiation, which generally lasts several weeks or months, as a convenient approach to generate disease models where aging is relevant, which is the case of AD and other neurodegenerative diseases (Jothi and Kulka, 2024; Pitrez et al., 2024). Moreover, careful manipulation of cellular reprogramming and differentiation protocols can enhance the recapitulation of disease-specific phenotypes *in vitro* (Jothi and Kulka, 2024). This approach not only extends cell culture age but also mimics natural developmental transitions that leave epigenetic marks required for proper function at maturity (Basu and Tiwari, 2021; Olynik and Rastegar, 2012). For comparison, transcription factor-based methods, like neurogenin 2 (NGN2), convert stem cells into the desired phenotype in several days, resulting in younger cultures that sometimes fail to recapitulate signatures characteristic of neurodegenerative disorders (Rothstein et al., 2023).

As chemical differentiation protocols may introduce significant variability in cell cultures, to enhance protocol consistency and throughput, we optimized key experimental variables and introduced thorough quality controls. Our protocol allows for large-scale cultures with reproducible molecular features. This method generates well-characterized batches of millions of NPCs with high expression levels of EMX1, and low expression levels of both DLX5, NKX2.1. This profile of transcription factor expression is characteristic of progenitors that give rise to glutamatergic neurons of the upper layers of the cortex, one of the most affected cell types in AD (Strano *et al*., 2020). Moreover, these NPCs can be cryopreserved without compromising their viability and phenotypic identity. To guarantee robustness, we validated the protocol in two different hiPSCs clones of two different lines. We further streamlined the terminal differentiation of NPCs to cortical neurons in 96-well plate formats, making these cell systems suitable for high throughput disease modelling and drug testing.

To develop an AD model, we refined available terminal differentiation protocols (Shi *et al*., 2012a; Strano *et al*., 2020) by implementing two key modifications. First, we introduced a top-up medium change regime (see Methods). From days *in vitro* 14 (DIV14) onwards, instead of completely replacing the culture medium, we supplemented (topped up) each well with fresh medium containing concentrated factors every 2-3 days. This approach was specifically designed to prevent the removal of soluble Aβ from the supernatant, thereby facilitating its aggregation. Secondly, we incorporated a chronic treatment with TNF-α at a final concentration of 20 ng/mL, applied from DIV14 to DIV41. This extended exposure to TNF-α was intended to mimic aspects of AD chronic neuroinflammation and potentially accelerate or enhance Aβ aggregation in our neuronal cultures.

Chronic TNFα stress induced Aβ puncta accumulation in the neurites and axonal projections of APP^Swe^ cortical neurons, confirmed with a panel of antibodies directed against APP, APP/Aβ or only Aβ isoforms. Notably, the fact that we observed that APP specific antibodies also differentially stained clusters in neurites between TNFα and vehicle treated cells suggest that TNFα-driven pathways may not only interfere with APP processing, increasing the Aβ levels, but may affect APP expression or degradation (Blasko et al., 1999; Liao et al., 2004; Plantone et al., 2023; Zhao et al., 2011). Moreover, formation of APP clusters in cell membranes has been reported to be linked to amyloidogenic processing (Schneider et al., 2008).

TNFα-induced Aβ accumulation also correlated with synaptic dysfunction and a hypermetabolic profile. Our results show that TNFα treatment disrupted synaptic structure shown by decreased SYN1 and PSD-95 co-localization, reduced neurogranin and HOMER1 levels, and triggered an increase of asynchronous electrical activity. Notably, these changes occurred after Aβ deposition peaks, suggesting a link between amyloid pathology and synaptic dysfunction.

The impact of TNFα in synapses has been extensively studied under physiological conditions, where this cytokine at non-pathological concentrations seems to play a key role in modulating neuronal function by regulating synaptic plasticity and long-term potentiation (LTP) processes (Heir and Stellwagen, 2020). For instance, TNFα is reported to increase the expression levels of AMPA receptors to strengthen glutamatergic synapses (Pickering et al., 2005). By contrast, pathological levels of TNFα in the brain of AD patients has been correlated with cognitive decline (Serafini *et al*., 2024). This evidence led to the development of clinical candidates, such as etanercept, directed to block TNFα to enhance cognitive performance of patients (Torres-Acosta et al., 2020). Despite strong evidence linking high TNFα levels with synaptic dysfunction, no human cellular models, like the one in this study, demonstrate that chronic exposure to TNFα disrupts synapses by altering the levels and distribution of key pre-synaptic and post-synaptic markers. By filling that gap, we provide a valuable tool for understanding and targeting TNFα-related synaptic dysfunction in AD.

Lastly, mitochondrial metabolism analysis revealed TNFα treatment also led to neuronal hypermetabolism. TNFα treated neurons demonstrated a higher oxygen consumption rate, revealed by a higher basal respiration and maximal respiration. Also, significantly higher proton leak levels and lower mitochondrial dependency for oxygen consumption in TNFα-treated neurons suggested that mitochondrial impairment was more pronounced under a neuroinflammatory stress. Notably, these phenotypes precede the increase in Aβ puncta levels, indicating that these alterations may be an early pathological event in our model, as it is reported in patients (Bakhtiari *et al*., 2024). Our metabolism results are consistent with previous reports showing that TNFα triggers very quick effects on mitochondrial dysfunction in neurons (Doll et al., 2015).

## Conclusions

We reported a robust platform to generate large numbers of well-characterized cortical NPCs and neurons in miniaturised, multi-well culture, formats. By applying this protocol, we generated APP^Swe^ neurons and subjected them to a chronic treatment with TNFα to model AD. These neurons replicate three early AD phenotypes: Aβ deposition in neuronal projections, synaptic dysfunction, and hypermetabolism. We anticipate that this platform will enable high throughput applications, and serve as a translational tool for drug discovery.

## Materials and Methods

### Study design

This study presents a protocol for generating hiPSC-derived cortical neurons to model key hallmarks of AD, using two hiPSC lines: one carrying the Swedish mutation (APP^Swe^) and a wild-type isogenic control, with two clones from each line. Quality controls are implemented at multiple stages. For hiPSC cultures, we perform morphology checks, maintaining <80% confluence, and assess expression of pluripotency markers (OCT4, SOX2, SSEA4) using immunofluorescence and flow cytometry. We further validate pluripotency by performing trilineage differentiation tests, checking for mesoderm (NCAM, Brachyury), endoderm (CXCR4, SOX17), and ectoderm (NESTIN, PAX6) markers. hiPSCs are differentiated to cortical NPCs, which are characterised by immunocytochemistry. Three different protocols are tested and scaled-up to generate batches of millions of NPCs for high-throughput applications. Only NPC batches meeting the following expression thresholds are kept for neuronal differentiation (>70% for EMX1, <50% for DLX5, and <20% for NKX2.1). Good quality NPCs should give rise to neuronal cultures enriched in TBR1 and with minimal contamination of astrocytes (GFAP) and oligodendrocytes (O4). Cell identity is assessed by immunocytochemistry. NPCs that pass the quality controls are used to develop AD models. To model AD pathology, we use APP^Swe^ neurons subject them to a chronic treatment with (TNF-α) at 20 ng/mL from DIV14 to DIV41, along with a top-up medium change regime to facilitate Aβ aggregation. We assess multiple AD-related phenotypes using various techniques: immunocytochemistry with multiple antibodies to detect Aβ aggregation in the neuronal body and neurites, expression and co-localization analysis of synaptic markers (SYN1/PSD-95, neurogranin and HOMER1), microelectrode array (MEA) recordings for electrophysiological activity, and oxygen consumption rate (OCR) measurements for mitochondrial metabolism. All cultures and assays are performed in microplates, aiming to establish a platform suitable for both AD pathology studies and high-throughput drug screening applications.

### hiPSC culture

Two clones from two hiPSCs lines were used in this study: BIONi10-Ca, BIONi10-Cb (RRID:CVCL_1E68) and BIONi10-C-38 APP KM670/671ML (RRID:CVCL_UR27), clones *a* and *b* generated by Bioneer (BION, Denmark). The BIONi10-C line is the parental line or isogenic control of BIONi10-C-38. The BIONi10-C was derived from dermis fibroblast cells (#0000208364, Lonza) of a male individual, of age ranging 15-19 years old and of Black/African-American ethnicity. The individual had no diagnosed disease at collection time. Further details about the line can be found in the hPSCreg database (https://hpscreg.eu/).

Both lines were validated, tested for karyotype, and maintained *Mycoplasma* free. hiPSCs were cultured in T75/T175 flasks coated with GelTrex™ (#A141320, Gibco) using mTeSR Plus medium (#100-0276, StemCell Technologies). For passaging, hiPSC colonies were treated with ReLeSR™ (#100-0484, StemCell Technologies) for 1 min. After removing the passaging reagent, the cells were incubated at 37 °C for 2 to 3 min. The colonies were then gently washed out of the flask with mTeSR, collected in a 50-mL sterile tube, and allowed to settle for 5 min at room temperature. The supernatant was carefully aspirated, and the colonies were gently resuspended in mTeSR before being transferred to freshly-coated flasks, minimising the pipetting steps. iPSCs were subcultured keep 1:4 to 1:6 splitting ratios.

Whenever appropriate, iPSCs were cryopreserved preserving colony integrity in the CryoStor® CS10 solution (#07959, StemCell Technologies). Approximately a 70-80% confluent T75 flask was cryopreserved in 5-7 cryovials (1 mL/each). After thawing, hiPSCs were cultured for a minimum of two weeks, and 4-6 passages, before use in experimental applications.

### Differentiation of hiPSCs to ectoderm, mesoderm and endoderm

hiPSCs were differentiated into the three germ layers (ectoderm, mesoderm, and endoderm) as follows. At day 0, the hiPSC colonies were dissociated into single cells using ACCUTASE™ (#07920, STEMCELL Technologies). The cells were then plated to a density of 10^5^ cells/cm^2^ on Matrigel-coated plates (#354277, Corning) and incubated overnight at 37 °C in mTeSR™1 medium to allow cell adherence. Trilineage differentiation was performed using the STEMdiff ki (#354277) using the respective ectoderm, endoderm or endoderm media. For ectoderm differentiation, the cells were cultured for 7 days with medium changes every other day. For mesoderm and/or ectoderm, the cells were culture for 5 days, with the medium being replaced every other day.

### Flow cytometry

Flow cytometry analysis was performed using FIX & PERM® Cell Fixation and Cell Permeabilization Kit (GAS003, Invitrogen). hiPSCs or lineage progenitors were harvested with accutase and resuspended PBS with 2% fetal bovine serum (FBS) (FACS buffer). After harvesting, the cells were passed through a 40 µm cell strainer to generate a single-cell suspension. Cells were centrifuged at 300 g for 5 min and the supernatant was discarded. The cell pellet was resuspended in 100 µL of Fixation Medium (Component A from the FIX & PERM® Kit) and incubated at room temperature for 15 min. After fixation, cells were centrifuged at 300 g for 5 minutes and the fixation medium was removed. The cell pellet was resuspended in 100 µL of Permeabilization Medium (Component B from the FIX & PERM® Kit) and incubated with primary antibodies specific for lineage markers (e.g., PAX6 for ectoderm, Brachyury for mesoderm, SOX17 for endoderm) for 30 min at room temperature. Cells were washed twice with FACS buffer by centrifuging at 300 g for 5 min and resuspending in 500 µL of FACS buffer each time. After the final wash, cells were resuspended in 100 µL of FACS buffer. The suspension was incubated with fluorophore-conjugated secondary antibodies for 30 min at room temperature in the dark. Cells were washed twice with FACS buffer as described above and resuspended in 300 µL of FACS buffer for analysis. Labelled cells were analysed on a flow cytometer to assess the expression of lineage-specific markers. This protocol ensures efficient fixation, permeabilization, and staining of hiPSCs for flow cytometry, allowing for the accurate assessment of differentiation status.

### Generation of cortical progenitors from hiPSCs

hiPSCs were differentiated to cortical progenitor cells as follows. At d0, hiPSC colonies were dissociated into single cells using StemPro Accutase (#A1110501, Gibco). These cells were then plated to full confluency in GelTrex™-coated flasks and incubated overnight at 37 °C in mTeSR with the addition of a Rho kinase (ROCK) inhibitor prior to neural induction. The neural maintenance medium (NMM) was prepared by combining N-2 and B-27-containing media in a 1:1 ratio. N-2 medium consisted of DMEM/F-12 GlutaMAX, 1X N-2 (#17502048, Gibco), 5 µg/mL of human insulin (#I9278, Sigma), 100 µM nonessential amino acids (#11140050, Gibco), and 100 µM 2-mercaptoethanol. B-27 medium was composed of Neurobasal (#21103049, Gibco) and 1X B-27 (#17504044, Gibco). To form a densely packed neuroepithelial layer, the hiPSC medium was replaced daily with NMM supplemented with 10 µM SB431542 and 100 nM LDN-193189 for 10 to 12 days. Afterwards, neuroepithelium was treated with PluriSTEM™ Dispase-II Solution (#SCM133, Sigma Aldrich) and manually fragmented into clumps of 100 to 200 µm. These clumps were replated in GelTrex™-coated flasks for rosette induction and incubated overnight in non-supplemented NMM to ensure proper attachment. From approximately day 11 to 13, the clump morphology was closely monitored. When rosettes appeared (typically around days 15 to 16), cells were treated with 1 µg/mL of CHIR-99021 (CHIR) and 20 ng/mL of FGF2 in NMM for four consecutive days. Subsequently, cells received daily treatment with 1 µg/mL of CHIR until their cryopreservation on days 19 to 22.

### Generation and maintenance of APP^Swe^ cortical neurons stressed with TNFα

Cortical progenitor cells were thawed to GelTrex™-coated 6-well plates in NMM. Cells were fed with fresh medium every day for 3 days prior replating. For replating, progenitors were dissociated with accutase. Cell suspension was centrifuged at 300 g for 5 min at room temperature and resuspended in NMM supplemented with the ROCK inhibitor and penicillin–streptomycin (#1514012, Gibco). Cells were plated in CellCarrier Ultra 96-well plates (PerkinElmer) coated with 0.002% poly-L-ornithine (#P4957, Sigma Aldrich) and 10 µg/mL of laminin (#L2020, Sigma Aldrich), at a density of 90k to 100k cells/well. At DIV1, 4, 7 and 11, 50% of the medium was replaced by fresh NMM supplemented with 20 ng/mL of rhBDNF (#248-BDB, R&D Systems), 20 ng/mL of rhGDNF (#212-GD, R&D Systems), 400 µM of L-ascorbic acid (#A4544, Sigma), 1 mM of cAMP (#D0627, Sigma), 2X of CultureOne (#A3320201, Gibco), and 20 U/mL of penicillin-streptomycin (Pen/Strep). Additionally, cells were treated with a final concentration of 10 µM DAPT (#2634/10, Tocris) at DIV1 and DIV4. At DIV14, a top up medium change regime was applied. For that, 50 µl of medium were removed per well and replaced with 20 µL of supplemented medium. After DIV14, 20 µL of supplemented medium were added per well every three days until the completion of the experiment. Details on medium composition and top up periodicity can be found in **Table S1**.

### Immunocytochemistry

Cells were fixed at room temperature for 10 min using a solution of 4% PFA (#28906, Thermo Scientific) diluted in D-PBS (+/+) (#14040141, Gibco). After fixing, the cells were washed three times with D-PBS (+/+), then permeabilized with 0.1% v/v Triton X-100 (#85111, Thermo Scientific) in D-PBS (+/+) for 30 min at room temperature. Next, the cells were treated with immunofluorescence blocking buffer (#12411, Cell Signaling Technology) for 1 h at room temperature. The blocking solution was removed, and the cells were incubated with the appropriate primary antibody solution overnight at 4 °C. Following four washes with D-PBS (+/+), the cells were incubated with Alexa Fluor-conjugated secondary antibodies for 1 h at room temperature in the dark. Both primary and secondary antibodies were diluted in immunofluorescence blocking solution. The cells were washed four more times with D-PBS (+/+) before staining the nuclei with Hoechst 33342 (#H3570, Thermo Fisher) at a 1:1000 dilution in D-PBS (+/+). A detailed list of the primary and secondary antibodies used in this study can be found in **Table S2**.

### Electrophysiology

Differentiation of NPCsinto neurons on MEA plates and recording of their electrophysiological activity involved several key steps. Initially, cortical NPCs were thawed on GelTrex™-coated 6-well plates in NMM at full confluency. The medium was refreshed daily for three days before transferring the cells to MEA plates. CytoView MEA 96 well plates (Axion Biosystems) were coated with 80 µL of 0.07% PEI solution per well, prepared in 1X Borate Buffer, and incubated at 37 °C for one hour. The plates were then washed twice with over 260 µL of sterile D-PBS per well and once with the same amount of sterile water. Care was taken to avoid scratching the electrode surface with pipette tips. After aspirating the water, the plates were left to air-dry overnight in a sterile biological safety cabinet. NPCs were collected as single cells with the use of accutase and counted. A total of 14.4 million cells were centrifuged and resuspended in 1.2 mL of dotting medium (100 µg/mL of L2020 in 1X N2B27 with 10 µM Rock inhibitor) to achieve a concentration of 12 million cells/mL, ensuring no bubbles formed. 10 µL of the cell suspension was carefully placed in the centre of each well, covering the electrodes. To prevent evaporation, sterile warm water was added to the surrounding plate reservoirs, and the plate was incubated at 37 °C for 1 h. Subsequently, 200 µL of 1X N2B27 with ROCK inhibitor was gently added to each well to avoid dislodging the adhered NPCs. NPCs were then differentiated into mature cortical glutamatergic neurons and treated with TNFα at a final concentration of 20 ng/mL every 2 or 3 days. For recording spontaneous electrical network activity, a 20-min equilibration period was followed by a 10-min measurement. Recordings were conducted using the Axion Integrated Study (AxIS) software in a controlled environment at 37 °C and 5% CO2. Electrical activity was measured with a gain of 1000x and a sampling frequency of 12.5 kHz. A Butterworth band-pass filter, ranging from 100 to 3000 Hz, was applied before spike detection. The AxIS adaptive spike detector set the threshold at six times the root mean square (RMS) noise on each electrode. An electrode was deemed active if it showed a spike rate of at least five spikes per min. These parameters were used to assess and characterize the spontaneous electrical network activity in the experimental setup. Each recording was saved as .spk files using the AxIS software. Network bursts were identified using the Axion Neural Metric Too.

### Assessing Mitochondrial Function Using the Seahorse XF Mito Stress Test Kit

A Seahorse XF Mito Stress Test Kit (Agilent Technologies, Cat. #103015-100) was utilized to measure mitochondrial function. For the assay setup, NPCs were treated with accutase to generate single cells and plated into the Seahorse XF96 Cell Culture Microplate at a density of 40k to 80k cells per well in NMM supplemented with Rock inhibitor. NPCs were differentiated to neurons and treated with the stressor TNFα as reported in previous methods sections (details in Table S1). On the day of the assay, the medium was replaced with fresh Seahorse XF DMEM Medium, and the plate was placed in a non-CO2 incubator at 37 °C for 1 h to equilibrate. The Seahorse XF Calibrant Solution (Agilent Technologies, Cat. #100840-000) was loaded into the Seahorse XF Sensor Cartridge and hydrated overnight. During the assay, the Mito Stress Test reagents, including oligomycin, FCCP, and a mix of rotenone and antimycin A, were sequentially injected into each well to measure key parameters of mitochondrial function, such as basal respiration, ATP production, maximal respiration, and spare respiratory capacity. Data were collected and analysed using the Seahorse XF software.

