## Supplementary Information for "TNF⍺-driven Aβ aggregation, synaptic dysfunction and hypermetabolism in human iPSC-derived cortical neurons"

**Table S1. Medium composition and factor concentration used for top-up medium change regime.**

| DIV | Volume to remove (μL) | Volume to add (μL) | Final volume per well (μL) | Medium composition |
|---|---|---|---|---|
| DIV14 | 50 | 20 | 70 | NMM, 35 ng/mL BDNF, 35 ng/mL GDNF, AA 700 μM, cAMP 1.75 mM, Pen/Strep 3.5X (-/+ TNFα at 70 ng/mL) |
| DIV18 | 0 | 20 | 90 | NMM, 45 ng/mL BDNF, 45 ng/mL GDNF, AA 900 μM, cAMP 2.25 mM, Pen/Strep 4.5X (-/+ TNFα at 90 ng/mL) |
| DIV21 | 0 | 20 | 110 | NMM, 55 ng/mL BDNF, 55 ng/mL GDNF, AA 1100 μM, cAMP 2.75 mM, Pen/Strep 5.5X (-/+ TNFα at 110 ng/mL) |
| DIV25 | 0 | 20 | 130 | NMM, 65 ng/mL BDNF, 65 ng/mL GDNF, AA 1300μM, cAMP 3.25 mM, Pen/Strep 6.5X (-/+ TNFα at 130 ng/mL) |
| DIV28 | 0 | 20 | 150 | NMM, 75 ng/mL BDNF, 75 ng/mL GDNF, AA 1500μM, cAMP 3.75 mM, Pen/Strep 7.5X (-/+ TNFα at 150 ng/mL) |
| DIV32 | 0 | 20 | 170 | NMM, 85 ng/mL BDNF, 85 ng/mL GDNF, AA 1700μM, cAMP 4.35 mM, Pen/Strep 8.5X (-/+ TNFα at 170 ng/mL) |
| DIV35 | 0 | 20 | 190 | NMM, 95 ng/mL BDNF, 95 ng/mL GDNF, AA 1900μM, cAMP 4.75 mM, Pen/Strep 9.5X (-/+ TNFα at 190 ng/mL) |
| DIV39 | 0 | 20 | 210 | NMM, 105 ng/mL BDNF, 105 ng/mL GDNF, AA 2100μM, cAMP 5.25 mM, Pen/Strep 10.5X (-/+ TNFα at 210 ng/mL) |
| DIV42 | 0 | 20 | 230 | NMM, 115 ng/mL BDNF, 115 ng/mL GDNF, AA 2300μM, cAMP 5.75 mM, Pen/Strep 11.5X (-/+ TNFα at 230 ng/mL) |

### Table S2. List of antibodies used in this study.

| LIST OF ANTIBODIES USED IN THIS STUDY | | | |
|---|---|---|---|
| TARGET | HOST, ISOTYPE, CLONE | CATALOG NUMBER | WORKING DILUTION |
| OCT4 | Rabbit, IgG | 2750, Cell Signalling Tech | ICC 1:500, IF 1:100 |
| SOX2 | Rabbit, IgG | 3579, Cell Signalling Tech | ICC 1:500, IF 1:100 |
| SSEA4 | Mouse, IgG3 | 4755, Cell Signalling Tech | ICC 1:500, IF 1:100 |
| Brachyury | Rabbit, IgG | ab20680, Abcam | ICC 1:500, IF 1:100 |
| NCAM | Mouse, IgG1, HCD56 | 60021, StemCell Tech | ICC 1:500, IF 1:100 |
| CXCR4 | Rabbit, IgG | ab124824, Abcam | ICC 1:500, IF 1:100 |
| SOX17 | Mouse, IgG1 | ab84990, Abcam | ICC 1:500, IF 1:100 |
| NESTIN | Mouse, IgG1, 196908 | ab6320, Abcam | 1:1000 |
| OTX1/2 | Rabbit, IgG | AB9566-I, Sigma Aldrich | 1:250 |
| PAX6 | Rabbit, IgG | 42-6600, Invitrogen | 1:500 |
| EMX1 | Rabbit, IgG | PA5-35373, ThermoFisher | 1:50 |
| DLX5 | Rabbit, IgG, EPR4488 | ab109737, Abcam | 1:1000 |
| NKX2.1/TTF1 | Rabbit, IgG, EP1584Y | ab76013, Abcam | 1:500 |
| Ki67 | Mouse, IgG1 κ | 550609, BD Biosciences | 1:600 |
| TBR1 | Rabbit, IgG | ab31940, Abcam | 1:1000 |
| GFAP | Rabbit, serum | AB5804, Sigma Aldrich | 1:1000 |
| O4 | Mouse, IgM, O4 | MAB1326, Bio-Techne | 1:500 |
| Anti-APP | Mouse, IgG1, kappa, LN27 | 13-0200, Invitrogen | 1:500 |
| Anti-APP/Aβ | Mouse, IgG2aκ, W02 | MABN10, Sigma Aldirch | 1:500 |
| Aβ | Mouse IgG1, κ, 3A1 | 847501, BioLegend | 1:50 |
| Aβ42 | Mouse IgG1, κ, 12F4 | 805509, BioLegend | 1:50 |
| TUJ1 | Rabbit, IgG | ab52623, Abcam | 1:1000 |
| SYN1 | Mouse, IgG1, 7H10G6 | MA5-31919, Invitrogen | 1:1000 |
| PSD-95 | Rabbit, IgG | ab18258, Abcam | 1:1000 |
| Neurogranin | Rabbit, IgG | PA5-76392, Invitrogen | 5 µg/mL |
| Homer1 | Rabbit, IgG | ab184955, Abcam | 1:500 |

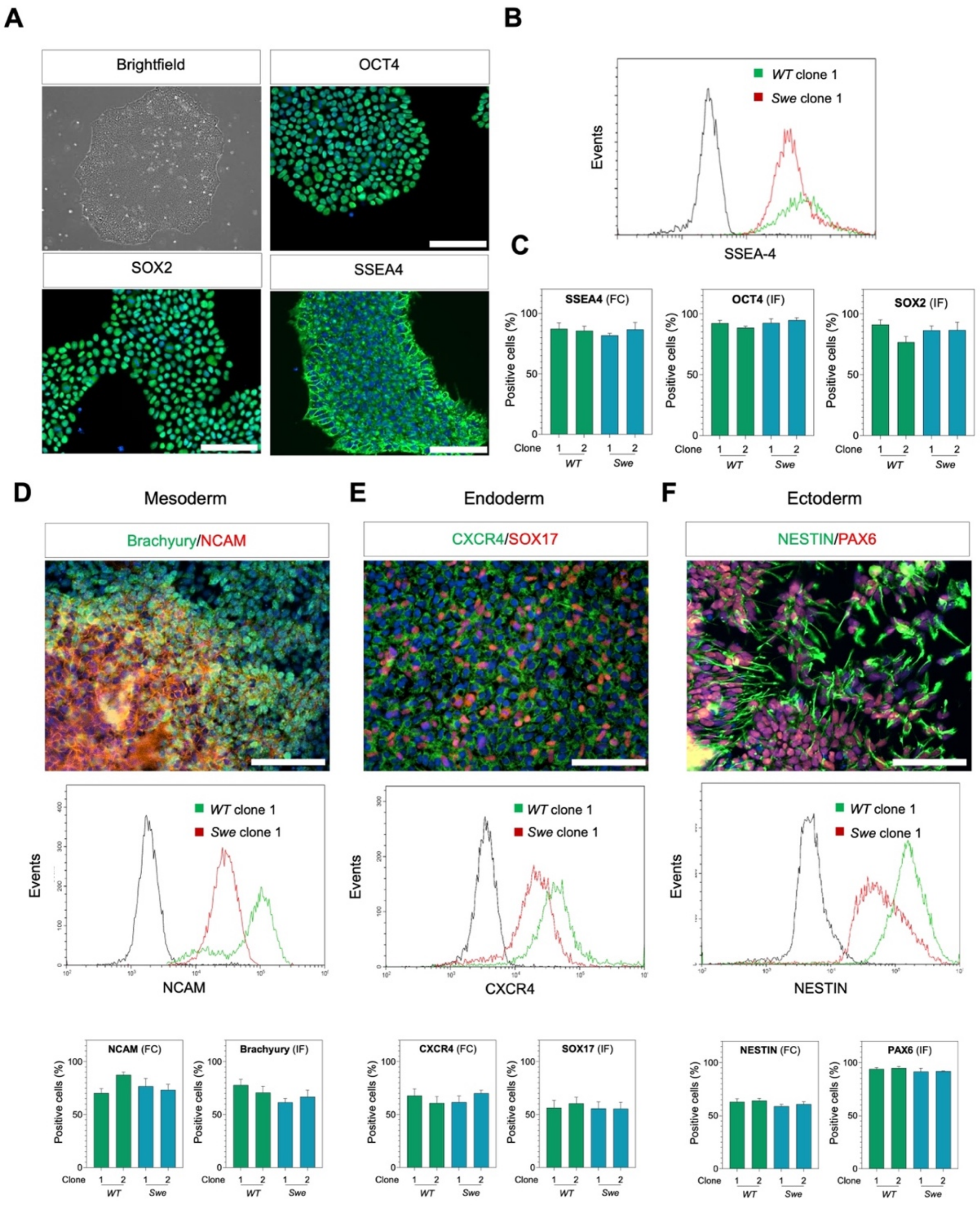


**Figure S1. Quality controls of wild-type and APP$^{Swe}$ mutant hiPSC lines. (A)** Representative phase contrast and ICC images of wild-type (*WT*) and APP$^{Swe}$ (*Swe*) mutant hiPSC colonies. **(B)** Representative flow cytometry histogram comparing the expression of the pluripotency marker SSEA4 in wild-type (green) and APP$^{Swe}$ (red) hiPSC cultures. Isotype control (black) is shown for reference. **(C)** Quantification of pluripotency markers across genotypes and clones. OCT4 and SOX2 were quantified via immunofluorescence (IF) as in A by scoring positive green nuclei relative to total viable DAPI-positive nuclei. SSEA4 was quantified via flow cytometry (FC) as in B relative to total viable cells. (**D-F**) Trilineage differentiation test. hiPSCs were differentiated to either mesoderm (D), endoderm (E) and ectoderm (F) progenitors to assess their pluripotency. Specific markers of each lineage were quantified (bottom panels) via IF (top panels) or FC (mid panels), that is, Bracryury and NCAM for Mesoderm, CXCR4 and SOX17 for Endoderm and NESTIN and PAX6 for Ectoderm. In **C-F,** triplicate data were normalized to total viable cells and are presented as mean ±SE. Scale bar = 100 μm.

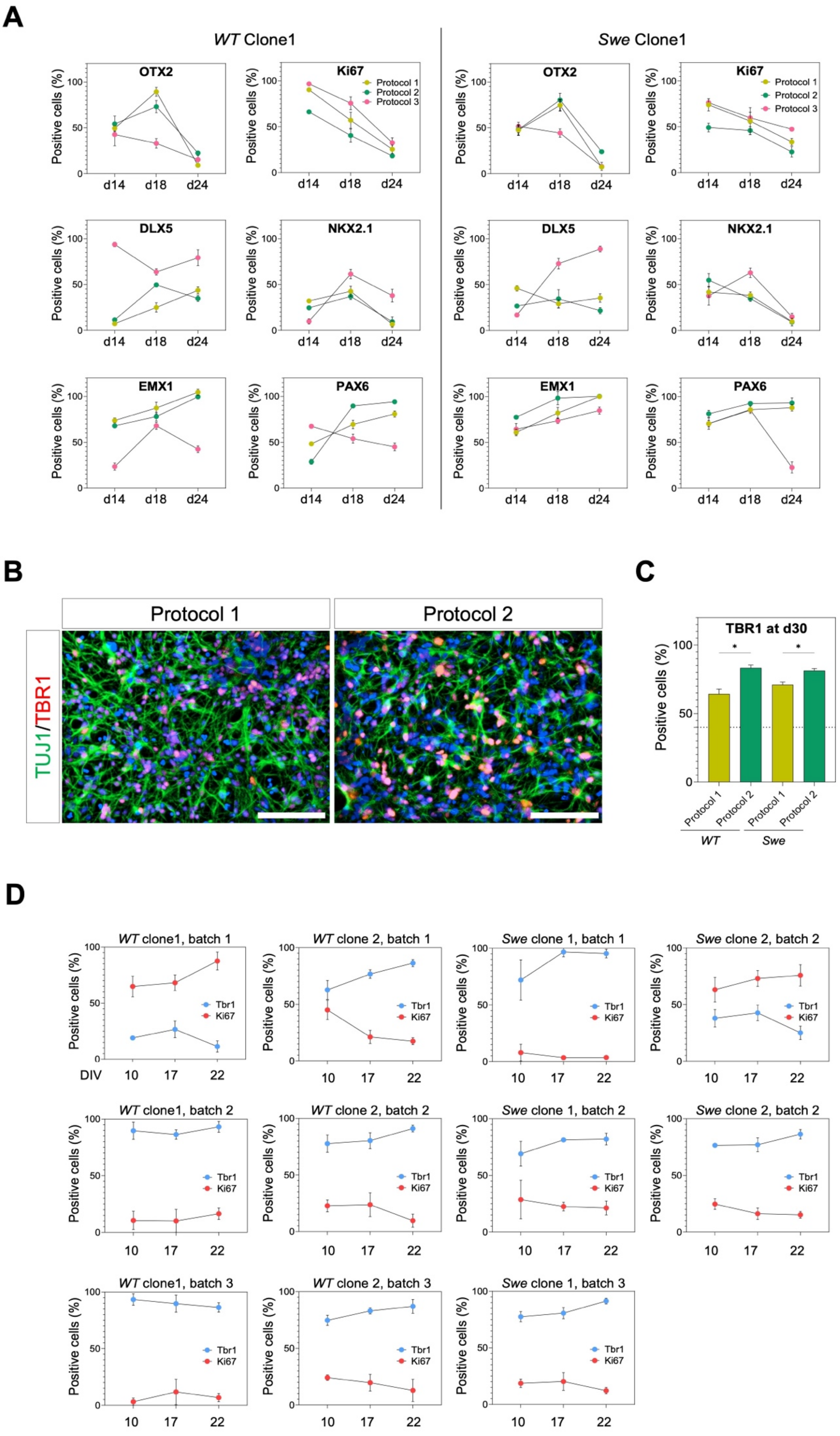
A
WT Clone1
Swe Clone1
OTX2
Ki67
DLX5
NKX2.1
EMX1
PAX6
Protocol 1
Protocol 2
Protocol 3
Positive cells (%)
d14
d18
d24
B
Protocol 1
Protocol 2
TUJ1/TBR1
C
TBR1 at d30
Positive cells (%)
Protocol 1
Protocol 2
WT
Swe
D
WT clone1, batch 1
WT clone 2, batch 1
Swe clone 1, batch 1
Swe clone 2, batch 2
WT clone1, batch 2
WT clone 2, batch 2
Swe clone 1, batch 2
Swe clone 2, batch 2
WT clone1, batch 3
WT clone 2, batch 3
Swe clone 1, batch 3
Tbr1
Ki67
DIV
10
17
22

**Figure S2. Quality controls of NPC and mature neurons from wild-type and APP$^{Swe}$ mutant lines.** **(A)** Quantification of differentiation markers in wild-type clone 1 and APP$^{Swe}$ clone 1 NPCs at different differentiation stages (d14, d18, d24). NPCs were generated using three protocols: Protocol 1 involved double SMAD inhibition followed by FGF2 induction, Protocol 2 introduced the use of CHIR during rosette development (d12-d17) and Protocol 3 implemented the use of the SHH inhibitor cyclopamine from d3-d9. The percentage of cells positive for OTX2, Ki67, DLX5, NKX2.1, EMX1, and PAX6 was measured via immunofluorescence relative to total healthy cells. **(B)** Representative immunofluorescence images of neurons differentiated using Protocol 1 and Protocol 2. Cells were stained for TUJ1 (green) and TBR1 (red). **(C)** Bar graph comparing the percentage of TBR1-positive cells at day 30 of differentiation (from hiPSC stage) across different protocols and genotypes. **(D)** Longitudinal IF-based quantification of Tbr1 (neuron marker) and Ki67 (proliferation marker) in wild-type and APP$^{Swe}$ clones across different NPC batches. In **A, B** and **D** triplicate data were normalized to total healthy cells and are presented as mean ±SE. In **C**. Two-way ANOVA was carried out to identify differences between differentiation Protocols across genotypes. Significance levels are as follows: (*) $P < 0.05$. Scale bar = 100 µm.
